# Abnormal reward prediction error signalling in antipsychotic naïve individuals with first episode psychosis or clinical risk for psychosis

**DOI:** 10.1101/214437

**Authors:** Anna O Ermakova, Franziska Knolle, Azucena Justicia, Edward T Bullmore, Peter B Jones, Trevor W Robbins, Paul C Fletcher, Graham K Murray

**Affiliations:** Department of Psychiatry, University of Cambridge; Behavioural and Clinical Neuroscience Institute, University of Cambridge; Cambridgeshire and Peterborough NHS Foundation Trust; Department of Psychology, University of Cambridge; Institute of Metabolic Science, University of Cambridge

**Keywords:** Substantia nigra, DLPFC, reinforcement learning, midbrain, predictive, ARMS

## Abstract

Ongoing research suggests preliminary, though not entirely consistent, evidence of neural abnormalities in signalling prediction errors in schizophrenia. Supporting theories suggest mechanistic links between the disruption of these processes and the generation of psychotic symptoms. However, it is not known at what stage in psychosis these impairments in prediction error signalling develop. One major confound in prior studies is the use of medicated patients with strongly varying disease durations. Our study aims to investigate the involvement of the meso-cortico-striatal circuitry during reward prediction error signalling in the earliest stages of psychosis. We studied patients with first episode psychosis (FEP) and help-seeking individuals at risk for psychosis due to subthreshold prodromal psychotic symptoms. Patients with either FEP (n = 14), or at-risk for developing psychosis (n= 30), and healthy volunteers (n = 39) performed a reinforcement learning task during fMRI scanning. ANOVA revealed significant (p<0.05 family-wise error corrected) prediction error signalling differences between groups in the dopaminergic midbrain and right middle frontal gyrus (dorsolateral prefrontal cortex, DLPFC). Patients with FEP showed disrupted reward prediction error signalling compared to controls in both regions. At-risk patients showed intermediate activation in the midbrain that significantly differed from controls and from FEP patients, but DLPFC activation that did not differ from controls. Our study confirms that patients with FEP have abnormal meso-cortical signalling of reward prediction errors, whilst reward prediction error dysfunction in the at-risk patients appears to show a more nuanced pattern of activation with a degree of midbrain impairment but preserved cortical function.

## Introduction

The cognitive basis of psychotic symptoms remains unknown, but abnormalities in the processing of prediction error (the mismatch between expected and actual outcome) have been proposed to contribute to the development of psychotic symptoms ^1,2^. Prediction errors can lead to allocation of attention and attribution of salience to stimuli, and drive subsequent learning ^3–5^. Faulty prediction error signalling could lead to several maladaptive psychological processes that have been proposed to contribute to the generation of psychotic symptoms: aberrant assignment of attention and motivational importance to innocuous stimuli, and disrupted associative learning leading to the formation of irrelevant associations and eventually delusions ^6–12^.

Several studies have attempted to examine the neural basis of prediction error abnormalities in psychosis, and have documented blunted midbrain, striatal, and/or cortical encoding of reward prediction errors ^13–16^ and non-reward related prediction errors ^17^. Our previous work in psychosis patients has identified meso-cortico-striatal prediction error deficits, involving midbrain, striatum and frontal cortex, especially right lateral frontal cortex ^13,17^. However, a major complication in the interpretation of patient studies is that several studies are potentially confounded by having either all, or the majority of patients, taking antipsychotic medication, which has been shown to modulate brain reward processing in healthy individuals and patients ^18–20^. Given this, and the likely importance of dopaminergic dysfunction in the pathogenesis of psychosis ^21^, it is critical to investigate possible abnormalities during reward prediction error processing in unmedicated and antipsychotic naïve patient samples. To our knowledge, only two studies ^16,22^ have examined reward prediction error signalling in unmedicated, but not all antipsychotic naïve, samples of mixed first episode and chronic schizophrenia patients (average age 27 years, ^16^, average age 34 years ^22^). Whilst both of these studies document striatal reward prediction error abnormalities, neither study report abnormalities in the dopaminergic midbrain. This is of particular interest given the extensive evidence for the role of dopamine in both prediction error signalling ^23,24^ and the pathophysiology of psychosis.

Some studies of chronic medicated schizophrenia patients have shown intact prediction error associated brain signals ^15,25,26^. It is possible that these abnormalities may be more prominent early in the course of the illness, especially in unmedicated samples. Establishing the pathophysiological abnormalities at the very earliest stages of illness is likely to be critical for optimal treatment and preventative interventions. The onset of psychosis is usually preceded by a prodromal phase involving social, educational or occupational decline accompanied by prodromal symptoms such as suspiciousness or hallucinations without a delusional interpretation ^27^. Help-seeking patients with these features have been shown to have increased risk for developing psychotic illness, and have been termed to have at-risk mental states (ARMS) or be at “ultra-high clinical risk” of psychosis ^27^. The prodromal phase may offer a critical period for intervention to improve long-term prognosis. The study of at-risk patients has proved a useful paradigm to investigate some of the earliest pathophysiological changes in schizophrenia and related illness ^28^. Brain prediction error signalling has not been examined in this group before, although there is some evidence for abnormal cortical and/or striatal processing of salience ^29,30^ or reward anticipation in at-risk patients ^31,32^. Given the theoretical importance of prediction error in learning and the pathogenesis of symptoms, we reasoned that it is key to examine brain prediction error signals in the earliest possible stages of psychosis. In this study, we set out to examine fMRI-correlates of reward prediction errors in a sample of patients with FEP and in at-risk patients, all naïve to antipsychotic medication, with a particular focus on the midbrain, striatum and right lateral frontal cortex.

A simple view of the continuum of psychosis is that at-risk patients with subthreshold symptoms will show similar pathology to clinical psychosis but of lesser severity ^33^ and there is evidence in support of this ^28^. We therefore hypothesized that patients with FEP would have abnormal prediction error activity in the dopaminergic midbrain, striatum and right lateral frontal cortex compared to controls, and that at-risk patients would have brain prediction error activation patterns intermediate between FEP patients and controls.

## Methods

### Participants

The study was approved by the Cambridgeshire 3 National Health Service research ethics committee. Individuals with FEP (n=14, average 23.57 years, 7 female) or at-risk for psychosis individuals (n=30, average 22.03 years, 14 female) were recruited from the Cambridgeshire first episode psychosis service, CAMEO. Inclusion criteria were as follows: age 16–35 years, early psychosis as reflected by meeting either at-risk attenuated psychotic symptoms criteria or FEP criteria according to the Comprehensive Assessment of At Risk Mental States (CAARMS) ^27^. Patients with FEP were required to be within 1 year of first presentation to the clinical service for psychosis; all participants were required to be naïve to antipsychotic medication. Healthy volunteers (n=39, average 23.23 years, 20 female) without a history of psychiatric illness or brain injury were recruited as control subjects. All subjects had normal or corrected-to-normal vision and no contraindications to MRI-scanning. None of the participants had a recreational drug or alcohol dependence. Healthy volunteers did not report any personal or family history of neurological, psychiatric or medical disorders, and were matched to patients regarding age, gender, handedness and maternal level of education. There were no significant differences between groups in age, sex, handedness, or recreational drug use (Supplementary Table 1). We used the Culture Fair matrices test, which is a very general IQ measure. There was a significant difference between groups in IQ. The three groups were not intended to be fully matched on IQ, as cognitive impairment is common in psychosis, and the task is not intellectually taxing. However, the groups were matched in maternal education, which indicates intellectual potential was matched. Even though the groups slightly differed in IQ, we did not find group differences in the performance. Furthermore, we did not find significant correlations between performance and IQ in any of the trial types. Therefore, we are confident that the group difference observed in our sample is due to psychiatric differences rather than differences in IQ. On average, the patients had predominantly positive psychotic symptoms and low levels of negative symptoms. FEP had significantly more severe symptoms than the at-risk group. Written informed consent was supplied by all participants.

### fMRI reward task

During the fMRI-scan, participants performed an instrumental discrimination learning task (Figure 1) involving monetary gains that required them to choose between two abstract visual stimuli (fractal pictures) ^13,18,34,35^. Three different pairs of stimuli are used: reward, neutral and bivalent stimuli. Each stimulus has an assigned outcome probability, from which participants learn how to maximise monetary payoff. Participants however could only learn to reliably win money in the reward trials, as the outcome in the bivalent was arbitrarily assigned to one stimulus independently of the participant’s choice and the neutral trials did not have any monetary value. Full details on the task are presented in the Supplementary Material.

**Figure 1.**
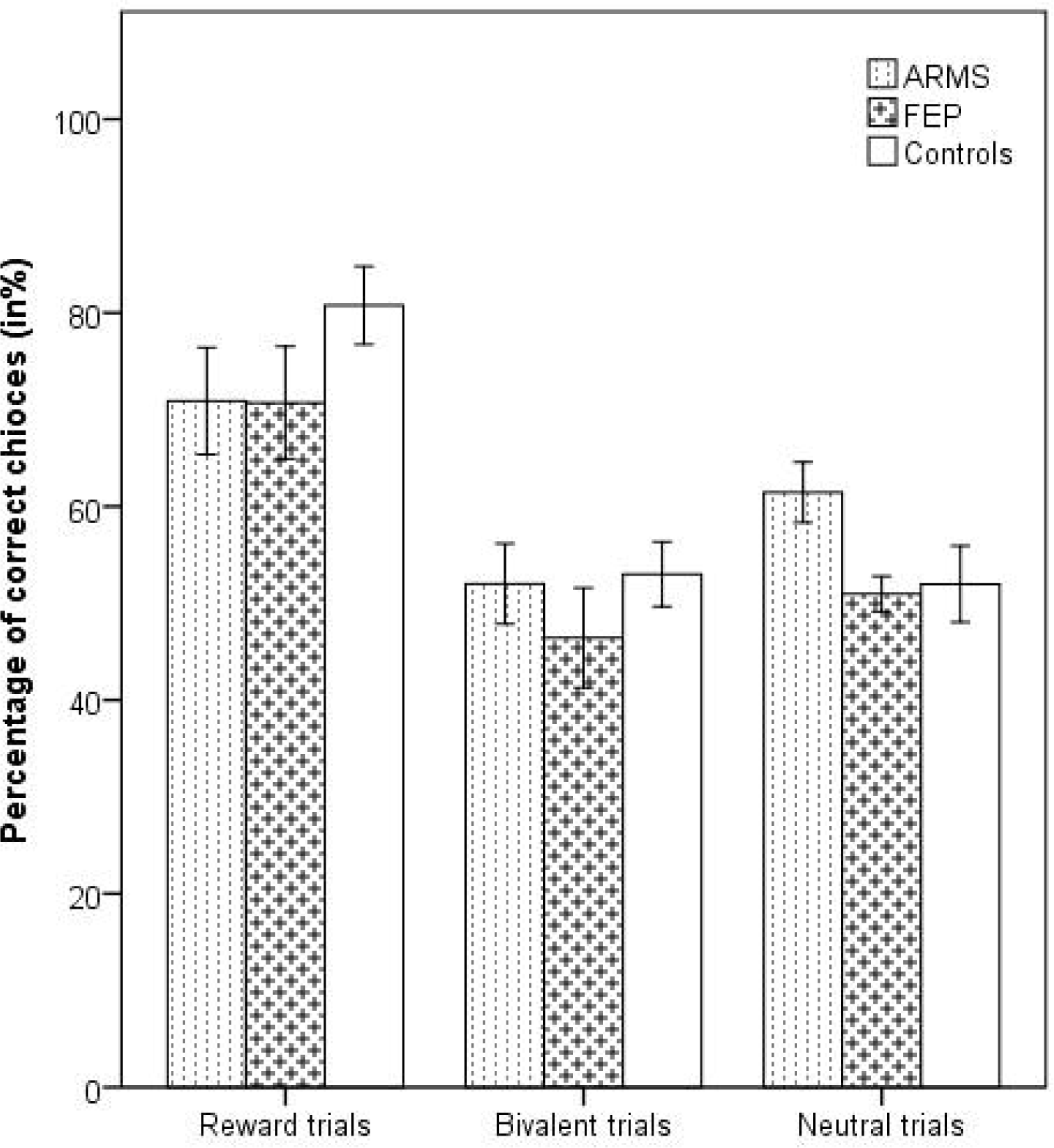
Behavioural Task. Panel A presents the three different trial types and feedback probabilities. Panel B presents the experimental task, including trial timing.

### Behavioural analysis

A 3x3 mixed-model analysis of variance (ANOVA, group x trial type) was used to investigate the group differences in reaction times and stimuli choices in the three types of trials (reward, bivalent and neutral). We examined the proportion of “correct” responses. Here the term “correct” means selecting the picture that leads to a high probability of getting £1 in the reward trial, and the picture with the higher probability of receiving the blue feedback picture in neutral trials, and is randomly assigned to choosing one of the pictures in bivalent trials (Figure 1, panel A). In the bivalent and neutral trials, the assignment of “correctness” is arbitrary, but assigning one stimulus in each category to be the “correct” stimulus allows examination of whether participants preferred one stimulus over another and the extent to which response patterns differed across trial types.

### fMRI data acquisition and analysis

Full details of our 3T scanning protocol and analysis methods are available in the Supplementary Material.

The seven explanatory variables (regressors) that we used were as follows: 1) onset of the bivalent cues; 2) onset of the neutral cues; 3) onset of the reward cues; 4) neutral outcome onsets (neutral feedback) during both neutral and reward trials; 5) winning outcome onset in the reward trials; 6) winning outcome onset in the bivalent trials; 7) loss outcome onset in bivalent trials. All regressors were modelled as 2 s events and convolved with a canonical double-gamma response function. We added temporal derivatives to the model to account for possible variation in the haemodynamic response function and we included motion parameters.

Our contrast of interest aimed at detecting activation associated with positive prediction error, and follows a contrast originally used by Seymour and colleagues who employed a similar paradigm in a healthy volunteer study in order to examine positive prediction error^34^. We contrasted winning £1 in bivalent trials versus winning £1 in reward trials; contrasting these identical outcomes in the context of different expectations represents a measure of positive prediction error. On the reward trials the reward is well predicted and elicits a low, yet still positive, prediction error; however, the outcome is unpredictable on bivalent trials and hence elicits a high positive prediction error ^34^. Therefore, contrasting the two events gives a measure of high versus low prediction error, and hence provides an assay of prediction error brain activation. This prediction error contrast has the advantage that it is perfectly balanced in terms of outcome value; hence it is unconfounded by reward outcome value or valence, which has been proposed to be a potential confound of alternative approaches, particularly in designs where reward prediction error and reward value are collinear ^36,37^.

For the group ANOVA analysis we used this prediction error contrast of interest as the outcome variable (FSL software calls outcome variables COPES: contrast of parameter estimates) and group as the predictor variable. We used permutation based statistics using the FSL tool randomise, utilising threshold-free-cluster enhancement, which enhances cluster-like structures but remains fundamentally a voxel-wise statistical testing method ^38^. We report results at p=0.05 or less, family-wise error corrected for multiple comparisons, using the variance smoothing option (3mm) as recommended for experiments with modest sample sizes, as is common in fMRI research ^39^. Our main analysis was based on a region of interest approach as follows. Our primary region of interest was the dopaminergic midbrain using the probabilistic atlas of Murty and colleagues ^40^, in which traditional anatomical segmentation was replicated using a seed-based functional connectivity approach and which provides a mask that includes substantia nigra and ventral tegmental area. The probabilistic map used to assess midbrain activation has been reliably used in a number of studies ^e.g.41,42^. In our two secondary regions of interest, we investigated the associative and limbic striatum (using a single hand-drawn mask, encompassing both associative and limbic striatum, based on operational criteria ^43,44^), and the right lateral frontal cortex (utilising a sphere, 10mm, centred at x=50, y30=, z=28, based on our previous work ^17^. Supplementary figure 1 shows our primary and secondary regions of interest. Whilst ANOVA will show whether the groups deviate from each other, it does not show the direction of effects. We therefore planned that for any voxels deemed significant in the ANOVAs, we would proceed to planned paired group comparisons, again using FSL randomise in order to test our hypothesis that on the contrast of interest, the following pattern would be seen: controls > at-risk > FEP). For each individual, we also extracted contrast values (contrast of parameter estimates, or COPEs in FSL) from voxels in which significant group differences were found, and calculated cluster averages; the extracted values were used for correlations with symptoms (within group), for bar charts of group differences and, as a supplement to our randomise paired group tests, to further test our hypothesis of controls > at-risk > FEP.

In support of this initial analysis and results, we also ran an additional analysis with a general linear model where a prediction error regressor was determined by a computational Q-learning model as we and others have used previously ^13,18,35^,^45^ (methods and results reported in Supplementary Material).

## Results

### Behaviour results: Choices

On analysis of “correct” choices across trial type and group, there was a main effect of trial type (F=20.93, df=2,160, p<0.001), but no effect of group (F=1.10, df=2,80, p=0.34) or group by trial type interaction (F=1.53, df=4,160, p=0.20; see Figure 2). The probability of choosing the “correct” stimulus increased in the course of the experiment (for a display of learning curves for each condition see Supplementary Figure 2 A-C.). Bonferroni corrected post-hoc tests showed that participants chose the “correct” stimulus more frequently in reward trials than in neutral (p<0.001), or bivalent (p<0.001), with no difference between neutral and bivalent trials (p=0.7). Participants learned to choose the picture that most often led to winning £1 (high-probability stimulus: “correct” response) in the majority of reward trials (mean percentage of “correct” choices 75%) whilst performance on other trials was similar to chance (neutral trials 55%, bivalent trials 52%).

**Figure 2.**
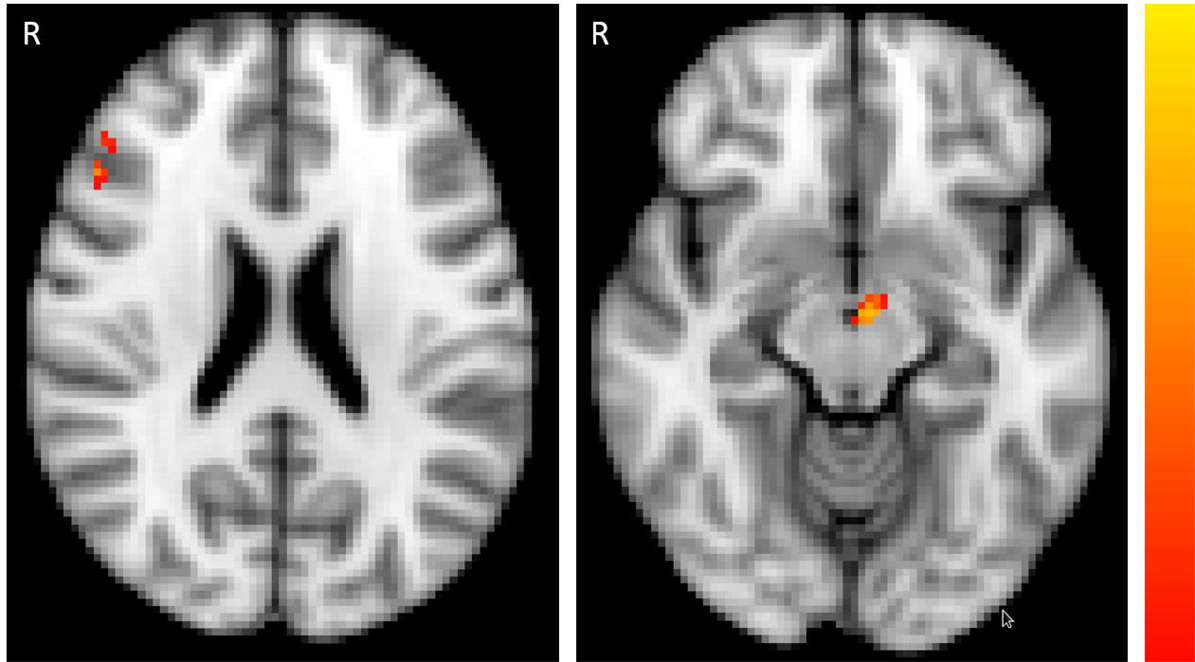
Trial performance: Percentage of “correct” (high-likelihood) choices stratified by trial type and participant group. Error bars are ±1 SE.

In order to investigate whether participants used the feedback to make an informed decision, further demonstrating engagement in the task, we conducted a win-stay lose-shift analysis (Supplementary Figure 3). We conducted a repeated measure 2 (win stay, lose shift) x 3 (reward, bivalent, neural) analysis across groups. Across all trial types and groups, we found a significantly higher stay probability after a rewarded (in the case of neutral: colour matching) trial compared to lose-shifting on unrewarded (non-matching) trials (win stay: 69.01% ±1.62, lose shift: 43.69% ±1.70; F=75.56, df=1,80, p<0.001). There was a significant trial type effect (F=6.0, df=2, p=0.003), as well as win stay/lose shift by group interaction (F=8.40, df=1,2, p<0.001), a win stay/lose shift by trial type interaction (F=32.79, df=1,2, p<0.001), and a marginally significant trial type by group interaction (F=2.26, df=2,2, p=0.065).

In our planned group comparisons (Supplementary Table 2 and Supplementary Figure 3), we found that controls had a significantly higher probability for a win stay behaviour than FEP patients on reward and neutral trial types (reward trials: p=0.027; neutral trials: p=0.012) and marginally on bivalent trials (p=0.062). Controls and at-risk patients were similarly likely to repeat the same response after a win. At-risk patients had a significantly higher probability of win stay behaviour in neutral trials compared to FEP (p=0.013), but the two patient groups did not differ on the other trial types. FEP patients had a higher probability for a lose shift behaviour compared to controls and at-risk patients in both reward and neutral trials (reward trials: FEP>Controls: p=0.001, FEP> at-risk: p=0.008; neutral trials: FEP>Controls: p=0.001, FEP> at-risk: p=0.045). Controls and at-risk patients were similarly likely to shift after a loss.

### Behaviour results: Reaction times

We analysed reaction times trial type and group. We found a significant main effect of trial type (F=4.473, df=2,160, p=0.01), but no effect of group (F=0.32, df=2,80, p=0.73) or group by trial type interaction (F=1.28, df=4,160, p=0.28). The Bonferroni corrected post-hoc tests revealed that participants independent of group reacted significantly (p<0.001) faster to reward trials (1122.83ms ±33.31) than to bivalent trials (1358.99ms ±41.49), and similarly (p=0.14) to neutral trials (1329.30ms ±97.87).

### Prediction error imaging results: ANOVA across three groups

We conducted second level (group level) ANOVAs using FSL randomise across the three groups across the whole brain and in our region of interests. Our outcome measure presented here in the main manuscript text is the contrast value (in FSL termed contrast of parameter estimates, or COPE) of a bivalent trial win versus a reward trial win, which corresponds to positive reward prediction error; group is the predictor variable. Results from a related outcome variable – prediction error associated brain activity derived from the computationally modelled prediction error – are presented in the Supplementary Material, and are similar. On whole brain analysis, there were no group differences that passed our statistical threshold corrected for multiple comparisons.

### Primary region of interest results: prediction error imaging results in the dopaminergic midbrain

We found a significant family-wise-error corrected group difference in the primary region of interest, the dopaminergic midbrain (maximal difference at x=-4, y=-12, z=-12; t=3.45, p=0.013 FWE corrected, 39 voxels; Figure 3). For each of these 39 significant voxels, family-wise error correcting for multiple comparisons, we performed planned group comparisons between pairs of groups using randomise to test our hypothesis of controls > at risk patients > FEP patients. The results were consistent with the hypothesis (Figure 4): at-risk patients were intermediate and significantly differed from both FEP patients and controls (controls>at-risk patients maximal difference at x=0, y=-16, z=-8; t=2.86, p=0.033 FWE corrected, 3 voxels; at-risk>FEP patients, maximal difference at x=-4, y=-8, z=-12; t=3.25, p=0.007 FWE corrected, 26 voxels). There was a significant difference between controls and FEP patients (controls>FEP, maximal difference at x=0, y=-16, z=-8; t=4.63, p<0.001 FWE corrected, 39 voxels). Another way to follow-up the significant effect of group in the ANOVA to test our hypothesis of brain prediction error signal following the pattern controls > at risk patients > FEP patients, is by taking the prediction error contrast value average for all 39 voxels that were significant in the ANOVA, and conducting planned paired group comparisons. This analysis also revealed that at-risk patients (mean=0.66, SD=54.86) were intermediate between the FEP patients and the controls, having a significantly smaller contrast value than controls: controls>at-risk patients, p=0.034, one-tailed; but significantly greater than in FEP: at-risk>FEP patients, p=0.001, one-tailed. The mean contrast value in the controls (mean=24.44, SD=50.12) was significantly greater than in FEP (mean=-55.03, SD=50.70): controls>FEP, <0.001, one-tailed, confirming our hypothesis of controls > at risk patient > FEP patients.

**Figure 3.**
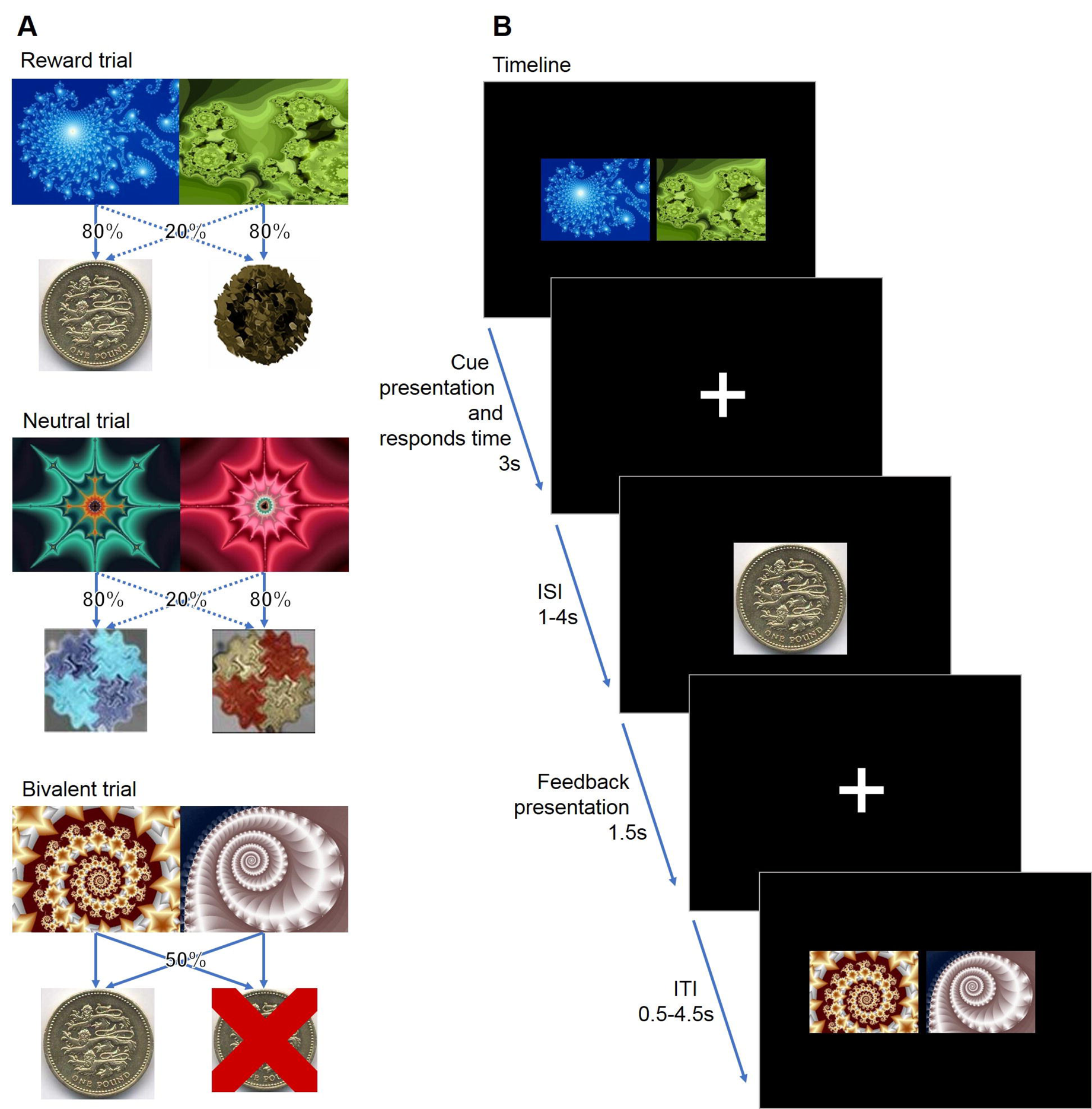
Group differences in region of interest analysis of activation associated with reward prediction error signal (p<0.05 FWE corrected) across the three groups (controls, first episode psychosis and at-risk patients) in the midbrain ventral tegmental area (right panel, z=-12), and right middle gyrus/dorsolateral prefrontal cortex (left panel, z=22). Colour bar depicts corrected voxel p-value from 0.001 (yellow) to 0.05 (red)).

**Figure 4.**
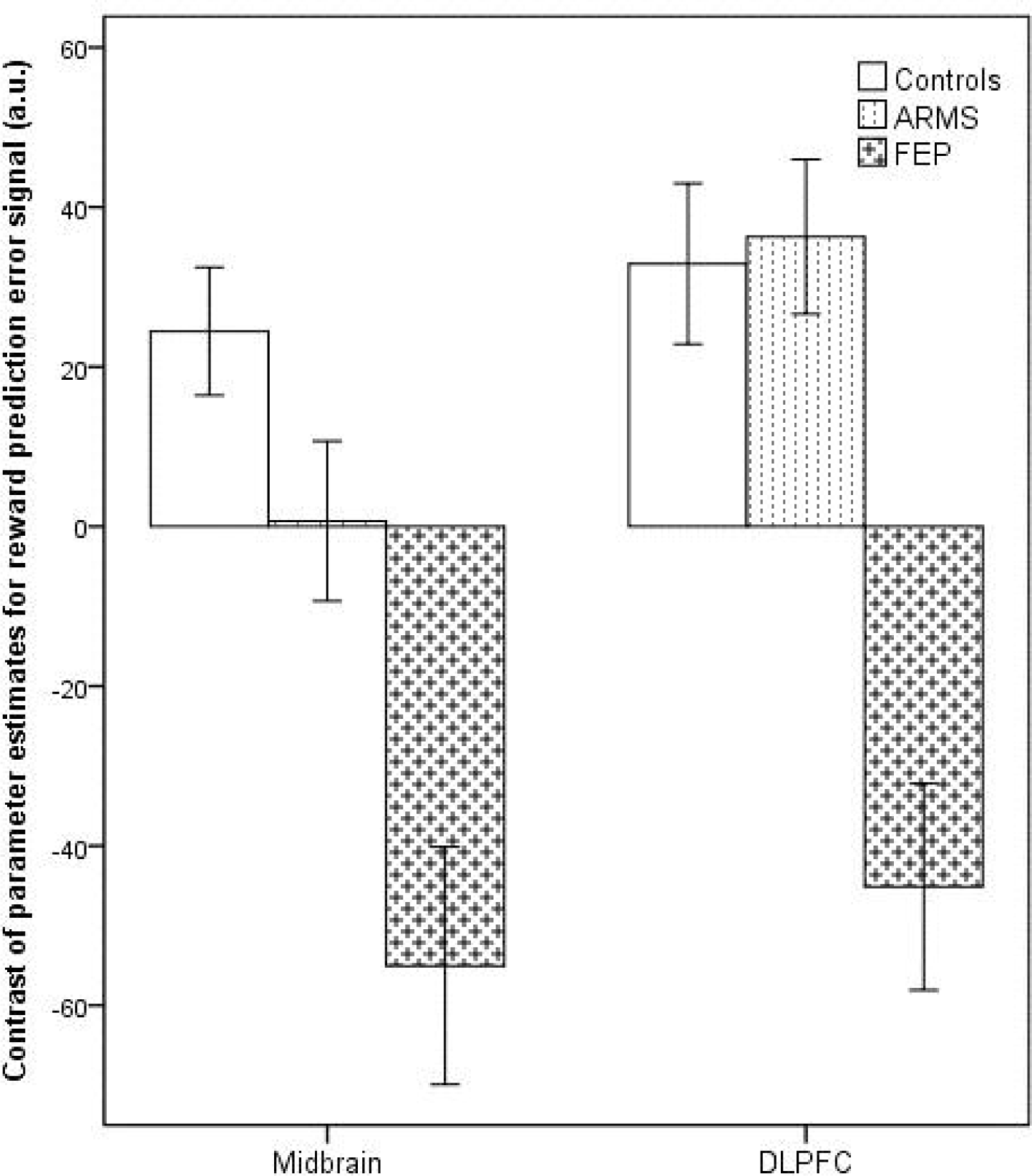
Bar chart shows the mean prediction error contrast values (termed contrast of parameter estimates, or COPEs in FSL) according to group, extracted from significant clusters determined by FSL randomise ANOVA results. The contrast values (COPEs) are derived from the contrast between Reward win and Bivalent win, which constitutes the reward prediction error. Error bars show ±1 SE; a.u. is arbitrary units.

### Secondary region of interest prediction error imaging results

No voxels passed our threshold on ANOVA in the associative-limbic striatum. There was a significant family-wise-error corrected group effect in the DLPFC (maximal difference at x=50, y=26, z=20; t=3.53, p=0.018 FWE corrected, 20 voxels; Figure 3). For each of these 20 significant voxels, family-wise error correcting for multiple comparisons, we performed planned two-group comparisons between using randomise to test our hypothesis of controls > at risk patients > FEP patients. The results were not consistent with the hypothesis (Figure 4). Whilst we found a significant difference between controls and FEP patients (controls>FEP, maximal difference at x=50, y=24, z=20; t=3.72, p<0.001 FWE corrected, 20 voxels), and between at-risk and FEP patients (at-risk>FEP, maximal difference at x=50, y=26, z=20; t=4.46, p<0.001 FWE corrected, 20 voxels), controls and at-risk patients did not differ significantly. We complemented the voxel based paired group comparisons with an analysis taking the prediction error contrast value average for all 20 voxels in the DLPFC that were significant in the ANOVA, and conducting planned paired group comparisons to test our hypothesis of brain prediction error signal following a pattern (controls > at risk patients > FEP patients). This analysis was consistent with the voxelwise paired group comparisons in that it did not support our hypothesis: the mean contrast values for these 20 voxels in controls (mean=32.90, SD=62.82) were significantly greater than in FEP (mean=-45.13, SD=48.40): controls>FEP, p<0.001, though not different from at-risk patients (mean=36.29, SD=53.03): controls>at-risk patients, p=0.81. Mean contrast values in at-risk patients were significantly greater than in FEP (p<0.001).

### Symptom correlations

There were no significant correlations between midbrain or DLPFC activation total CAARMS symptom severity in FEP (midbrain rho = 0.03, p=0.93; DLPFC rho=-0.27, p=0.34) or the at-risk group (midbrain rho = 0.18, p=0.33; DLPFC rho=0.233, p=0.21).

## Discussion

We show evidence of abnormal midbrain signalling of reward prediction errors in patients with FEP and at-risk for psychosis. In addition, we also report DLPFC abnormalities in patients with FEP, showing that abnormalities associated with prediction error processing in psychotic illness are not restricted to sub-cortical regions, consistent with previous results ^13,17,46,47^. These results are not secondary to antipsychotic medication, because people with current or previous prescriptions of these drugs were excluded. Therefore, our findings extend previous findings reporting abnormal reward prediction error signalling in the midbrain, striatum and cortex in a partly medicated sample of FEP patients (in which results held in a very small unmedicated subsample) ^13^, in medicated samples of schizophrenia patients ^14,15^, and in the cortex and striatum of unmedicated, but not antipsychotic naïve, samples of mixed FEP and chronic schizophrenia patients (mean age 27 years, ^16^, or mean age 34 years ^22^). We note that one previous study ^20^ tested an unmedicated, antipsychotic naïve, schizophrenia sample and also reported midbrain alterations in a reward associated task; this study, however, focussed on anticipation of reward and punishment rather than reward prediction error. To our knowledge, our study is, therefore, the first to document abnormal brain prediction error signals in the dopaminergic midbrain with early-stage psychosis in an entirely antipsychotic naïve sample.

Our study is also the first to examine reward prediction error signalling in patients at-risk for developing psychosis, and as such it will require replication before definitive conclusions can be drawn, especially as there were only very small areas of difference in the at risk patients. We found a mild abnormality in midbrain prediction error signalling in the at-risk group that was intermediate in severity between psychotic illness and controls. Whilst the prediction error signal in the dorsolateral prefrontal cortex was also disrupted in FEP patients, dorsolateral prefrontal cortex prediction error signalling was relatively spared in the at-risk group, contrary to our hypothesis. These findings were similar in both the contrast based prediction error analysis presented in the main manuscript text, and in the alternative computationally informed approach presented in the Supplementary Material.

Conflicting evidence exists as to whether the same pathological mechanisms are responsible for both prodromal (sub-threshold) and severe psychotic symptoms ^33^. Under the straightforward account of a dimensional theory of psychosis ^48^, the same pathology responsible for severe psychotic symptoms in schizophrenia should also be present (to a milder degree) in people with sub-threshold psychotic symptoms such as suspicions or mild hallucinations. Such an account would posit that people at-risk for psychosis due to the presence of sub-threshold psychotic symptoms would be characterised by a level of pathophysiological disruption that is of similar nature but lesser severity than the florid illness. The findings of the present study provide some support this theory for the dopaminergic midbrain, analogous to the pattern seen in a previous PET imaging study in the striatum ^28^. We do note, however, we did not find significant associations with symptoms scores, which would be expected by a dimensional account. Another possibility, however, is that there may be qualitative differences in the pathology of sub-threshold and severe psychosis. The currently still largely intact frontal prediction error signalling in the at-risk group speculatively may be a mechanism that helps prevent a mild symptom becoming a severe one (e.g., a suspicion becoming a delusion), although longitudinal studies would be required to test such a mechanistic hypothesis.

Our study was not designed to be sensitive to prediction error signalling in all cortical regions, and would be unlikely to be sensitive to auditory cortex predictive signals that have been implicated in the generation of auditory hallucinations ^49,50^. It would therefore be premature to conclude that all cortical prediction error signalling is intact in at-risk patients. We note that two previous studies found preliminary evidence (not corrected for multiple comparisons) of enhanced frontal activation anticipating a reward in clinical risk groups using the monetary incentive delay paradigm ^31,32^. Whilst these prior findings may appear to contrast with our results of intact activation in the dorsolateral prefrontal cortex in the at-risk group, the studies are consistent in showing more prominent subcortical reductions in signalling in the at-risk state with no evidence of cortical reductions.

The probabilistic map of the dopaminergic midbrain we used to assess midbrain activation ^40^ has been used in a number of prior studies ^41,42^. It combines the substantia nigra and the VTA; however, we did not seek to differentiate these structures. Differentiating activation amongst midbrain nuclei is generally challenging and confirmation of the precise anatomical abnormalities in psychosis could be facilitated with future developments in MRI technology such as the use of higher field strengths. We emphasize here that our results pertain to the region of the dopaminergic midbrain, acknowledging that the distinction between the substantia nigra and VTA is challenging in fMRI as well as that fMRI does not demonstrate the neurochemical origin of the signals observed. Our analysis of variance within a combined limbic and associative striatal region of interest did not demonstrate any voxels showing a significant group difference corrected for multiple comparisons. This could be due to a lack of power, with a small sample size in the FEP group and variable activation in the at-risk patients, or it could indicate areas of relatively spared function in some patients ^25,46,51,52^. The relatively small sample size, especially in the FEP group, is a limitation of the study.

In the analysis of choice performance and reaction time data we did not find any significant group differences. Win stay/lose shift analysis revealed that participants, independent of trial type (reward, bivalent or neutral), have a significantly higher probability to repeat the same response after a win than to shift after a loss, which is a clear indication for engagement and learning in a probabilistic learning task ^53,54^. The win stay probability was highest on reward trials, as expected, as these trials are the most predictable and consistently rewarding. However, also the win stay probability on bivalent trials shows that participants were engaging in the task and attempting to apply a learning strategy. The neutral trials show a lower probability rates and a less clear model learning behaviour. This, however, is not important for our analysis, as neutral trial imaging correlates are covariates of no interest. This analysis revealed subtle behavioural differences between the controls and the FEP patients: FEP patients have lower probabilities to repeat a response that lead to a win and a higher probability to shift after a loss, showing less stability in their decision making process.

An advantage of our study is that we use a paradigm that can generate assays of prediction error signalling either by a traditional cognitive subtraction fMRI approach or by a computational modelling fMRI approach (Supplementary Material) as previously applied by others and ourselves ^13,14,34,53^. The convergent results indicate that the findings are not secondary to particular modelling strategy supplied, and are not confounded by outcome valence, which is often highly collinear with prediction error in many reinforcement learning studies, which has been raised as a concern raised by some researchers previously ^36^.

In conclusion, we document midbrain and cortical abnormalities in prediction error signals in antipsychotic naïve FEP patients. The findings in our at-risk group suggest a more nuanced account of the pathogenesis of symptoms than that predicted by a simple continuum model of psychosis. In the at-risk group, there was midbrain evidence of dysfunction consistent with a dimensional account of psychosis, but relatively spared dorsolateral prefrontal cortical function in contrast to frank psychosis, arguably more consistent with a categorical model of psychiatric disorder ^54^. Further investigations into areas of continuity and discontinuity between the at-risk and frank psychosis patients, including longitudinal designs, may bring insights into factors critical in the pathogenesis of psychotic illness.

## Acknowledgments and funding

Supported by a MRC Clinician Scientist [G0701911] and an Isaac Newton Trust award to Dr Murray; by the University of Cambridge Behavioural and Clinical Neuroscience Institute, funded by a joint award from the Medical Research Council [G1000183] and Wellcome Trust [093875/Z/10/Z]; by awards from the Wellcome Trust [095692] and the Bernard Wolfe Health Neuroscience Fund to Dr Fletcher, and by awards from the Wellcome Trust Institutional Strategic Support Fund [097814/Z/11], and Cambridge NIHR Biomedical Research Centre. The authors are grateful for the help of clinical staff in CAMEO for help with participant recruitment and Wolfson Brain Imaging Centre staff for MRI data collection support.

## Author contribution

Dr Ermakova designed the study, collected data, performed behavioural and image analysis, interpreted results, and wrote the first draft of the manuscript. Dr Knolle designed and performed computational modelling, image analysis and model based image analysis, behavioural analyses, interpreted results, and substantially revised the manuscript. Ms Justicia designed the study, collected data, and performed behavioural analysis, and revised the mansucript. Dr Bullmore designed the study and revised the manuscript. Dr Jones designed the study and revised the manuscript. Dr Robbins designed the study and revised the manuscript. Dr Fletcher designed the study and revised the manuscript. Dr Murray had the idea for the study, designed the study, supervised and performed data acquisition, analysis, and interpretation, and wrote the manuscript. Dr Murray is the guarantor.

## References

1. Fletcher PC, Frith CD. Perceiving is believing: a Bayesian approach to explaining the positive symptoms of schizophrenia. Nat Rev Neurosci. 2009;10(1): 48–58. doi:10.1038/nrn2536.

2. Deserno L, Boehme R, Heinz A, Schlagenhauf F. Reinforcement learning and dopamine in schizophrenia: Dimensions of symptoms or specific features of a disease group? Front Psychiatry. 2013;4(DEC). doi:10.3389/fpsyt.2013.00172.

3. Dayan P, Kakade S, Montague PR. Learning and selective attention. Nat Neurosci. 2000;3 Suppl(november): 1218–1223. doi:10.1038/81504.

4. Pearce JM, Hall G. A Model for Pavlovian Learning: Variations in the Effectiveness of Conditioned But Not of Unconditioned Stimuli. Psychol Rev. 1980;87(6): 532–552. doi:10.1037/0033-295X.87.6.532.

5. Schultz W, Dickinson A. Neuronal coding of prediction errors. Annu Rev Neurosci. 2000;23: 473–500. doi:10.1146/annurev.neuro.23.1.473.

6. Robbins TW. Relationship between reward-enhancing and stereotypical effects of psychomotor stimulant drugs. Nature. 1976;(5581): 57–59. doi:10.1038/264057a0.

7. Miller R. Schizophrenic psychology, associative learning and the role of forebrain dopamine. Med Hypotheses. 1976;2(5): 203–211. doi:10.1016/0306-9877(76)90040-2.

8. Gray JA. Integrating schizophrenia. Schizophr Bull. 1998;24(2): 249–266.

9. Heinz A. Dopaminergic dysfunction in alcoholism and schizophrenia - Psychopathological and behavioral correlates. Eur Psychiatry. 2002;17(1): 9–16. 10.1016/S0924-9338(02)00628-4.

10. Kapur S. Psychosis as a state of aberrant salience: A framework linking biology, phenomenology, and pharmacology in schizophrenia. Am J Psychiatry. 2003;160(1): 13–23. doi:10.1176/appi.ajp.160.1.13.

11. Corlett PR, Frith CD, Fletcher PC. From drugs to deprivation: A Bayesian framework for understanding models of psychosis. Psychopharmacology (Berl). 2009;206(4): 515–530. doi:10.1007/s00213-009-1561-0.

12. Griffiths O, Langdon R, Le Pelley ME, Coltheart M. Delusions and prediction error: reexamining the behavioural evidence for disrupted error signalling in delusion formation. Cogn Neuropsychiatry. 2014;19(5): 439–467. doi:10.1080/13546805.2014.897601.

13. Murray GK, Corlett PR, Clark L, et al. Substantia nigra / ventral tegmental reward prediction error disruption in psychosis. Mol Psychiatry. 2008;13(3): 1–18. doi:10.1038/sj.mp.4002058.Substantia.

14. Morris RW, Vercammen A, Lenroot R, et al. Disambiguating ventral striatum fMRI-related BOLD signal during reward prediction in schizophrenia. Mol Psychiatry. 2012;17(3):235, 280–289. doi:10.1038/mp.2011.75.

15. Gradin VB, Kumar P, Waiter G, et al. Expected value and prediction error abnormalities in depression and schizophrenia. Brain. 2011;134(6): 1751–1764. doi:10.1093/brain/awr059.

16. Schlagenhauf F, Huys QJM, Deserno L, et al. Striatal dysfunction during reversal learning in unmedicated schizophrenia patients. Neuroimage. 2014;89: 171–180. doi:10.1016/j.neuroimage.2013.11.034.

17. Corlett PR, Murray GK, Honey GD, et al. Disrupted prediction-error signal in psychosis: Evidence for an associative account of delusions. Brain. 2007;130(9): 2387–2400. doi:10.1093/brain/awm173.

18. Pessiglione M, Seymour B, Flandin G, Dolan RJ, Frith CD. Dopamine-dependent prediction errors underpin reward-seeking behaviour in humans. Nature. 2006;442(7106): 1042–1045. doi:10.1038/nature05051.

19. Schlagenhauf F, Wüstenberg T, Schmack K, et al. Switching schizophrenia patients from typical neuroleptics to olanzapine: Effects on BOLD response during attention and working memory. Eur Neuropsychopharmacol. 2008;18(8): 589–599. doi:10.1016/j.euroneuro.2008.04.013.

20. Nielsen MO, Rostrup E, Wulff S, et al. Alterations of the brain reward system in antipsychotic naive schizophrenia patients. Biol Psychiatry. 2012;71(10): 898–905. doi:10.1016/j.biopsych.2012.02.007 [doi].

21. Laruelle M, Kegeles LS, Abi-Dargham A. Glutamate, Dopamine, and Schizophrenia from Pathophysiology to Treatment. In: Annals of the New York Academy of Sciences. Vol 1003.; 2003: 138–158. doi:10.1196/annals.1300.063.

22. Reinen JM, Van Snellenberg JX, Horga G, Abi-Dargham A, Daw ND, Shohamy D. Motivational Context Modulates Prediction Error Response in Schizophrenia. Schizophr Bull. 2016:sbw045. doi:10.1093/schbul/sbw045.

23. Schultz W, Dayan P, Montague PR. A neural substrate of prediction and reward. Science (80-). 1997;275(5306): 1593–1599.

24. Tian J, Huang R, Cohen JY, et al. Distributed and Mixed Information in Monosynaptic Inputs to Dopamine Neurons. Neuron. 2016. doi:10.1016/j.neuron.2016.08.018.

25. Culbreth AJ, Westbrook A, Xu Z, Barch DM, Waltz JA. Intact Ventral Striatal Prediction Error Signaling in Medicated Schizophrenia Patients. Biol Psychiatry Cogn Neurosci Neuroimaging. 2016. doi:10.1016/j.bpsc.2016.07.007.

26. Koch K, Schachtzabel C, Wagner G, et al. Altered activation in association with reward-related trial-and-error learning in patients with schizophrenia. Neuroimage. 2010;50(1): 223–232. doi:10.1016/j.neuroimage.2009.12.031.

27. Yung AR, Yuen HP, McGorry PD, et al. Mapping the onset of psychosis: The comprehensive assessment of at risk mental states. Schizophr Res. 2005;39(1): 964–971. doi:10.1016/S0920-9964(03)80090-7.

28. Howes OD, Montgomery AJ, Asselin MC, et al. Elevated striatal dopamine function linked to prodromal signs of schizophrenia. Arch Gen Psychiatry. 2009;66(1): 13–20. doi:10.1001/archgenpsychiatry.2008.514.

29. Smieskova R, Roiser JP, Chaddock CA, et al. Modulation of motivational salience processing during the early stages of psychosis. Schizophr Res. 2014;166(1–3): 17–23. doi:10.1016/j.schres.2015.04.036.

30. Roiser JP, Howes OD, Chaddock CA, Joyce EM, McGuire P. Neural and behavioral correlates of aberrant salience in individuals at risk for psychosis. Schizophr Bull. 2013;39(6): 1328–1336. doi:10.1093/schbul/sbs147.

31. Juckel G, Friedel E, Koslowski M, et al. Ventral striatal activation during reward processing in subjects with ultra-high risk for schizophrenia. Neuropsychobiology. 2012;66(1): 50–56. doi:10.1159/000337130.

32. Wotruba D, Heekeren K, Michels L, et al. Symptom dimensions are associated with reward processing in unrnedicated persons at risk for psychosis. Front Behav Neurosci. 2014;8. doi:10.3389/fnbeh.2014.00382.

33. Murray GK, Jones PB. Psychotic symptoms in young people without psychotic illness: Mechanisms and meaning. Br J Psychiatry. 2012;201(1): 4–6. doi:10.1192/bjp.bp.111.107789.

34. Seymour B, Daw N, Dayan P, Singer T, Dolan R. Differential encoding of losses and gains in the human striatum. J Neurosci. 2007;27(18): 4826–4831. doi:10.1523/JNEUROSCI.0400-07.2007.

35. Bernacer J, Corlett PR, Ramachandra P, et al. Methamphetamine-induced disruption of Frontostriatal reward learning signals: Relation to psychotic symptoms. Am J Psychiatry. 2013;170(11): 1326–1334. doi:10.1176/appi.ajp.2013.12070978.

36. Stenner M-P, Rutledge RB, Zaehle T, et al. No unified reward prediction error in local field potentials from the human nucleus accumbens: evidence from epilepsy patients. J Neurophysiol. 2015:jn.00260.2015. doi:10.1152/jn.00260.2015.

37. Rutledge RB, Skandali N, Dayan P, Dolan RJ. Dopaminergic Modulation of Decision Making and Subjective Well-Being. J Neurosci. 2015;35(27): 9811–9822. doi:10.1523/JNEUROSCI.0702-15.2015.

38. Winkler AM, Ridgway GR, Webster MA, Smith SM, Nichols TE. Permutation inference for the general linear model. Neuroimage. 2014;92: 381–397. doi:http://doi.org/10.1016/j.neuroimage.2014.01.060.

39. Nichols TE, Holmes AP. Nonparametric permutation tests for functional neuroimaging: A primer with examples. Hum Brain Mapp. 2002;15(1): 1–25. doi:10.1002/hbm.1058.

40. Murty VP, Shermohammed M, Smith D V, Carter RM, Huettel SA, Adcock RA. Resting state networks distinguish human ventral tegmental area from substantia nigra. Neuroimage. 2014;100: 580–589. doi:10.1016/j.neuroimage.2014.06.047.

41. MacInnes JJ, Dickerson KC, Chen N, Adcock RA. Cognitive Neurostimulation: Learning to Volitionally Sustain Ventral Tegmental Area Activation. Neuron. 2017;89(6):1331–1342. doi:10.1016/j.neuron.2016.02.002.

42. Bär K-J, de la Cruz F, Schumann A, et al. Functional connectivity and network analysis of midbrain and brainstem nuclei. Neuroimage. 2016;134: 53–63.

43. Mawlawi O, Martinez D, Slifstein M, et al. Imaging human mesolimbic dopamine transmission with positron emission tomography: I. Accuracy and precision of D(2) receptor parameter measurements in ventral striatum. J Cereb Blood Flow Metab. 2001;21(9): 1034–1057. doi:10.1097/00004647-200109000-00002.

44. Martinez D, Slifstein M, Broft A, et al. Imaging human mesolimbic dopamine transmission with positron emission tomography. Part II: amphetamine-induced dopamine release in the functional subdivisions of the striatum. J Cereb Blood Flow Metab. 2003;23(3): 285–300.

45. Erdeniz B, Rohe T, Done J, Seidler R. A simple solution for model comparison in bold imaging: the special case of reward prediction error and reward outcomes. Front Neurosci. 2013;7:116.

46. Murray GK, Corlett PR, Fletcher PC. The neural underpinnings of associative learning in health and psychosis: How can performance be preserved when brain responses are abnormal? Schizophr Bull. 2010;36(3): 465–471. doi:10.1093/schbul/sbq005.

47. Robbins TW. The case of frontostriatal dysfunction in schizophrenia. Schizophr Bull. 1990;16(3): 391–402. doi:10.1093/schbul/16.3.391.

48. Linscott RJ, van Os J. Systematic Reviews of Categorical Versus Continuum Models in Psychosis: Evidence for Discontinuous Subpopulations Underlying a Psychometric Continuum. Implications for DSM-V, DSM-VI, and DSM-VII. Annu Rev Clin Psychol. 2010;6(1): 391–419. doi:10.1146/annurev.clinpsy.032408.153506.

49. Horga G, Schatz KC, Abi-Dargham A, Peterson BS. Deficits in predictive coding underlie hallucinations in schizophrenia. J Neurosci. 2014;34(24): 8072–8082. doi:10.1523/JNEUROSCI.0200-14.2014.

50. Dowd EC, Frank MJ, Collins A, Gold JM, Barch DM. Probabilistic reinforcement learning in patients with schizophrenia: Relationships to anhedonia and avolition. Biol Psychiatry Cogn Neurosci Neuroimaging. 2016;1(5): 460–473.

51. Dowd EC, Barch DM. Pavlovian reward prediction and receipt in schizophrenia: Relationship to anhedonia. PLoS One. 2012;7(5). doi:10.1371/journal.pone.0035622.

52. Friston KJ, Price CJ, Fletcher P, Moore C, Frackowiak RS, Dolan RJ. The trouble with cognitive subtraction. Neuroimage. 1996;4(2): 97–104. doi:10.1006/nimg.1996.0033.

53. Frank MJ, Moustafa AA, Haughey HM, Curran T, Hutchison KE. Genetic triple dissociation reveals multiple roles for dopamine in reinforcement learning. Proc Natl Acad Sci. 2007;104(41): 16311–16316. doi:10.1073/pnas.0706111104.

54. Nowak M, Sigmund K. A strategy of win-stay, lose-shift that outperforms tit-for-tat in the Prisoner’s Dilemma game. Nature. 1993;364(6432): 56–58. doi:10.1038/364056a0.

55. Lawrie SM, Hall J, McIntosh AM, Owens DGC, Johnstone EC. The “continuum of psychosis”: Scientifically unproven and clinically impractical. Br J Psychiatry. 2010;197(6): 423–425. doi:10.1192/bjp.bp.109.072827.

